# PHYTOREMEDIATION OF REACTIVE TURQUOISE BLUE H5G USING *IN VITRO* CULTURES OF *SOLANUM VIRGINIANUM* (L.)

**DOI:** 10.1101/2022.07.26.501524

**Authors:** Dhanashree S. Patil, Swaroopa A. Patil

**Affiliations:** Department of Botany, Shivaji University Kolhapur

**Keywords:** Antioxidant, Biochemical, Phytoremediation, Reactive Turquoise Blue H5G, *Solanum virginianum*

## Abstract

*Solanum virginianum* (L.) belonging to family Solanaceae selected for decolourisation of Reactive Turquoise Blue H5G dye. *In vitro* grown cultures of *S. virginianum* were able to remove more than 50% dye concentration up to 110mg/l. comparative analysis of biochemical and antioxidant study showed more activity in treated plants as compared to untreated plants. The phytotoxicity study demonstrated the non-toxic nature of degraded metabolites. Use of such non-edible yet medicinal plant for phytoremediation is discussed.

## INTRODUCTION

Textile wet processing is considered as one of the worst industrial sectors in the terms of water consumption and water pollution. There are many textile industries in the country pollute the water bodies with the dyes in the effluent released. Hazardous chemicals are used for various processing in the textile industries as for scouring, desizing, bleaching, printing, dyeing and finishing. These chemicals include inorganic compounds, elements or polymers and organic products. More than 8000 chemical products are associated with the dying processes listed in colour Index (Society of Dyers and Colourists, 1976) whereas, 7×10^5^ metric tons of dyestuff is produced every year (Zollinger, 1987). Dyes are carcinogenic and mutagenic, toxic to the flora and fauna (Hu *et al*., 2009). These dyes include several structural varieties of dyes such as acidic, reactive, basic, disperse, azo, diazo, anthraquinone based and metal complex dyes. Colour, first contaminant of waste water has to be removed before releasing waste water into primary water bodies or on land. Apart from this as less than 1 ppm of the dye concentration is highly visible, it has magnified negative effect on aesthetics, transparency, gas solubility and overall the physico-chemical properties of water bodies (Banat *et al*., 1996). Dye containing effluent have a high chemical oxygen demand, biological oxygen demand, suspended solids and other toxic compounds. To mitigate these problems several physical, chemical and biological clean-up technologies are coming forward. The use of microbes (bioremediation) and plants (phytoremediation) for breaking down hazardous textile dyes is becoming a best choice to detoxify or render harmless environmental pollutants (Kagalkar *et al*. 2009; Patil *et al*. 2009). The costs of conventional physical and chemical treatment methods are highly expensive, have low efficacy, are labourers and though they assure decontamination, secondary pollutants may be second parallel problem hence, substitute for the method of decontamination is the need of an hour.

Compared to the physical and chemical technologies, bioremediation is an effective technique which is cost effective (Singh *et al*., 2008). Among the sources of bioremediation, the use of plants has opened up a new facet to clean up the contaminated soil. Researchers have realized that though the knowledge about detoxification and hyperaccumulation in plants is not well understood, plants are armored with amazing metabolic and absorption capabilities as well as transport systems that can uptake contaminants from soil and water and behave as a good remedy. Use of plants is safe, easy to operate and is less troublesome (Cunningham and Berti, 2000) because plant physiology and metabolism works in a way that mutations in the system will not spread as in case of microorganisms. Plants have extraordinary potential to concentrate and accumulate elements and compounds as well as organic contaminants and metabolize them in various molecules in their tissues. The plants have developed an advanced regulatory mechanism to co-ordinate effective metabolic activities (Salt *et al*., 1998). Dye decolorization ability of plants is mainly dependent upon the enzymes they consist of. Never the less the employment of plants for reclamation of highly polluted sites (water bodies or land) has come up. The utilization of non-edible plants for this purpose is rising day by day. Phytoremediation, a biological reclamation technique has proven to be alternative for physical and chemical techniques. Phytoremediation is environment friendly, less labourers, solar energy driven, toxin free, cost effective, vast applicable on various textile dyes, safer technology in terms of use and user and retrieval technology based on natural concept (UNEP, Undated; USEPA, 2000).

Reactive dyes are compounds of very simple chemical structure. Chemically these dyes consist azo compounds, anthraquinones and phthalocyanines. They possess high fixing property by simple dyeing methods, making covalent bridges with the fibre (cotton, wool or nylon), by the compatible hydroxyl group of cellulose. Reactive Turquoise Blue H5G (Reactive Blue 25) preoccupy excellent fastness properties, the dye is widely used in different textile industries for coloring linen, cotton, silk and other fibers. Textile industries uses different chemical containing dyes and released them in the form of millions of liters of untreated effluent. Reactive group of azo dyes is widely used in textile industrial processes.

*Solanum virginianum* L. (*Solanum surattense* Burm. f.; *Solanum xanthocarpum* Schrad. & H. Wendl.) is a diffuse and very prickly undershrub, belonging to the family Solanaceae. It can grow commonly in various regions of the world on sandy soils and is distributed throughout India. It is commonly called as yellow-berried nightshade in English, kantakari in Sanskrit, and nelagulla in Kannada. It is one of the members of “Dashamula” of Ayurveda. A wide range of phytochemicals such as phenolics, flavonoids, alkaloids, saponins, tannins, glycosides, fatty acids and amino acids have been reported previously from different parts of the plant (Dalavi and Patil, 2017). The plant is extensively used in various systems of medicine including Ayurveda. Plant serves as natural antioxidants. Antioxidants neutralizes the adverse effect of free radicals by inhibiting the initiation or propagation of oxidizing chain reactions generated by reactive free radicals like ROS (Reactive Oxygen Species), hydroxyl ion, superoxide, singlet oxygen and UV-B radiations before vital molecules are damaged. *Solanum virginianum*, a highly medicinal plant was selected for the study for degradation of Reactive Turquoise Blue H5G textile dye. The medicinal properties related to the biochemicals present in the plant and exploration of antioxidant property before and after treatment of dye was successfully carried out in the present investigation. The studies were confirmed with phytotoxicity analysis. Plant cell cultures offer several advantages to understand accurately the complexity of physiological and biochemical mechanism involved in the degradation of dyes, toxic chemicals and other pollutants (Doran, 2009). The present study indicates that the plant does not lose their potential even after treated with textile dyes and hence the plant can be successfully used in phytoremediation.

As mentioned above, plants are safe to handle. The selected plant shows many advantages such as, it is fast growing, has well developed root system, it is non-edible, non-hazardous and no special care is required. *S. virginianum* is used in traditional herbal medicines. Non-edible plant is chosen for phytoremediation because there is no risk of mixing experimental material in the routine food chain. In this regards *S. virginianum* would be an ideal system.

## 2. MATERIALS AND METHODS

### 2.1 Collection of plant material

Morphologically healthy fruits of *Solanum virginianum* (L.) were collected from Haripur, Sangli (16^0^ 50^’^ 52.2^”^ N 74^0^ 32^’^ 05.8^”^ E) (Maharashtra, India). The plant sample was submitted to SUK acronym (DSP001). The collected plants were maintained at botanical garden of Department of Botany, Shivaji University, Kolhapur.

### 2.2 *In vitro* regeneration of plant

Morphologically healthy seeds were selected as an explant. All the fruits were washed and surface sterilized with 0.1% HgCl2, followed by three successive washes of sterilized distilled water. Surface sterilization was carried out in laminar air flow. Fruits were blotted and dissected out. The seeds were then blotted on sterile tissue paper, dried and inoculated on ½ MS (Murashige and Skoog) (1962) basal medium aseptically. After germination they was subcultured on MS medium supplemented with different concentrations (0.5mg/l, 0.75mg/l and 1mg/l) of BAP (6 Benzylaminopurine).

### 2.3 Treatment of *in-vitro* plantlets with RTB H5G

Experiments were conducted to evaluate the efficiency of *S. virginianum* (L.) in removal/degradation of textile dyes Reactive Turquoise Blue H5G (RTB H5G). RTB H5G (synthetic dye) was procured from local market of Kolhapur. Different concentrations (10mg/l, 30mg/l, 50mg/l, 70mg/l, 90mg/l, 110mg/l, 130mg/l and 150mg/l) of dye were added to MS medium and a set of three Erlenmeyer flasks (100ml capacity) was prepared for each concentration. *In vitro* grown plantlets of *S. virginianum* were placed in all the treatment flasks. Additional two sets of three Erlenmeyer flasks each were prepared for abiotic control (Dye dissolved in distilled water instead of MS) and biotic control (MS Basal) all these flasks were incubated at 27±2 ºC for a period of six days under rotary shaking (110rpm). After day six aliquots (2ml from each flask separately) were collected and centrifuged at 8000rpm for 15min. The clear supernatant was used for measuring the residual dye (%) at 618nm, the absorbance readings were used for calculating degradation (decolourisation) of dye by the formula, % Decolourisation = (Initial absorbance - Final absorbance)/ Initial absorbance x100

The aliquots were collected after every 4-day period. Aseptic conditions were followed throughout the experiments.

### 2.4 Phytochemical analysis

After 21 days of decolourisation experiment, plants from all the different treatments were collected and plant extracts (1%) were prepared on fresh weight bases. Six different solvent systems, methanol, ethanol, acetone, chloroform, n-Hexane and distilled water were used for extraction.

#### 2.4.1 Quantification of total phenolic content (TPC)

The total phenolic contents of *S. virginianum* extracts were determined by using modified spectrophotometric Folin-Ciocalteu method (Wolfe *et al*., 2003). The reaction mixture was prepared by mixing an aliquot of extracts (0.125 ml) with Folin-Ciocalteu reagent (0.125 ml) and 1.25 ml of saturated Na_2_CO_3_ solution. Reaction mixture was further incubated for 90 min at room temperature and absorbance was measured at 760nm. The samples were prepared in triplicates for each analysis and the mean value of absorbance was recorded. Results were expressed as of mg gallic acid equivalents (GAE)/g fresh weight of samples of *S. virginianum*. All the experiments were expressed as mean ± SE of triplicate measurements.

#### 2.4.2 Quantification of total flavonoid content (TFC)

The total flavonoids were estimated by using modified colorimetric method (Luximon-Ramma *et al*., 2002). The reaction mixture had 1.5 ml of extract to 1.5 ml of 2% methanolic AlCl3. The mixture was incubated for 10 minutes at room temperature and absorbance was measured at 368nm against 2% AlCl3, which served as blank. The samples were prepared in triplicates for each analysis and the mean value of absorbance was obtained. The optical density (OD) measurements of samples were compared to standard curve of rutin and expressed as mg of rutin equivalent (RE)/100g fresh weight of plant *S. virginianum*. All the experiments were performed in triplicates and expressed as mean ± Standard Error (SE).

#### 2.4.3 Quantification of total alkaloid content (TAC)

Total alkaloid content of *S. virginianum* was assessed using phenanthroline method described by Singh *et al*., (2004). The assay mixture was prepared and incubated for 30 minutes in water bath maintained at 70±2^0^ C and the absorbance was measured at 510nm. Distilled water served as blank. The samples were prepared in triplicates for each analysis and the mean value of absorbance was recorded. The optical density (OD) measurements of samples were compared to standard curve of quercetin as mg of quercetin equivalent (QE)/100g fresh weight of *S. virginianum*. All the experiments were expressed as mean ± SE.

#### 2.4.4 Ferric reducing antioxidant power assay (FRAP)

The ferric ion reducing capacity was calculated by using assay described by Pulido *et al*. (2000). To 100 μl plant extract, 3 ml of FRAP reagent [300 mM sodium acetate buffer at pH 3.6, 10 mM of 2,4,6-Tripyridyl-S-triazine (TPTZ) solution and 20 mM FeCl3.6 H2O solution (10:1:1)] was added. The reaction mixture was incubated at 37°C for 15 min. The absorbance was measured at 595nm. The value of FRAP was expressed as milligrams of ascorbic acid equivalents per 100 gram of Fresh weight.

#### 2.4.5 2,2-diphenyl-1-picrylhydrazyl (DPPH) radical scavenging assay

The free radical scavenging activity of plant extract was measured (Aquino *et al*., 2001). Plant extract (25 μl) was mixed with 3 ml of DPPH methanolic solution (25 mM). The reaction mixture was incubated in dark at room temperature for 30 min. The absorbance was measured at 517nm against blank. Results were expressed as percentage of inhibition of the DPPH radical and percent antioxidant activity of plant extract was calculated using the following formula:

% DPPH Inhibition = [Control (abs) – Sample (abs)] ×100 /Control (abs)

#### 2.4.5 Ferrous ion chelating activity (FICA)

Ferrous ion chelating activity was measured by following method described by (Dinis *et al*., 1994). Assay mixture contained 0.1 ml of 2 mM FeCl2 and 0.3 ml of 5 mM ferrozine and mixed with 1 ml of plant extract. The mixture was incubated for 10 min at room temperature and absorbance was measured at 562nm spectrophotometrically. The ability of sample to chelate ferrous ion was calculated as the percent inhibition of Fe^+2^ to ferrozine complex. Percentage antioxidant activity of plant extract was calculated using the following formula:

% Ferrous ion inhibition = [Control (abs) – Sample (abs)] ×100 /Control (abs)

#### 2.4.6 Superoxide anion scavenging assay (SOAS)

Nitroblue tetrazolium (NBT) prepared in dimethyl sulfoxide (DMSO), was used for assaying SOAS. The reaction mixture was prepared by adding 0.1 ml of NBT (10 mg of NBT in 10 ml DMSO), 0.3 ml of plant extract and 1 ml of alkaline DMSO (1 ml of alkaline DMSO containing 0.1 ml of 5 mM NaOH and 0.9 ml of DMSO) was added to make final volume 1.4 ml and the absorbance was recorded at 560nm. DMSO solution was used as blank. Decrease in value of absorbance of the reaction mixture designate the increase in superoxide anion scavenging activity (Tiwari *et al*., 2017). The SOAS activity was calculated by using formula:

% SOAS inhibition = [Control (abs) – Sample (abs)] ×100 /Control (abs)

#### 2.4.7 Phosphomolybdenum reducing power assay (PMo)

Antioxidant capacity of the extracts was evaluated by phosphomolybdenum method according to the procedure described by Prieto *et al*. (1999). Plant extract (0.3 ml) was combined with 3 ml of reagent solution (0.6 M sulphuric acid, 28 mM sodium phosphate and 4 mM ammonium molybdate). The tubes containing the reaction solution were incubated at 95°C for 90 min. The absorbance of the solution was measured at 695nm using a UV-visible spectrophotometer against blank after cooling to room temperature. Methanol (0.3 ml) in the place of extract was used as the blank.

% Phosphomolybdenum inhibition = [Control (abs)–Sample (abs)] ×100/ Control (abs)

## 3. Phytotoxicity study

Ten seeds (*Vigna radiata*) of same size were placed in Petri plates (in triplicate) with blotting paper. Set of three Petri plates for each dye concentration (10mg/l, 30mg/l, 50mg/l, 70mg/l, 90mg/l, 110mg/l, 130mg/l and 150mg/l) and biotic control (MS basal) was used. Germination studies were carried out by watering the seeds with respective degraded solution every day. Seeds with radical (≥1mm) were considered germinated (Wu *et al*., 2007). The length of plumule (shoot), radical (root), fresh weight, rate of germination (%) was recorded after eight days. Germination percentage was calculated by the formula:

Germination (%) = (No. of seeds germinated / Total No. of seeds) X 100

## 4. RESULTS

### 4.1 *In vitro* regeneration of plant

Seed germination was achieved on ½ Murashige and Skoog basal medium (1962) (Table 1). Shoot multiplication, was carried out using *in vitro* nodal explants subcultured on MS medium supplemented with different concentrations of BAP (0.5mg/l, 0.75mg/l and 1mg/l). Highest shoot number (18.1±0.17, Table 2) was observed on MS medium supplemented with BAP (0.75mg/l).

### 4.1 Decolourisation of textile dye Reactive Turquoise Blue H5G (RTB H5G)

Dyes used in textile industries are with various chemical constituents. And hence it is a need to evaluate decolourisation efficiency of the plant for different concentration of dye. In the present study, screening of RTB H5G was carried out with eight different concentrations (10, 30,50,70,90,110,130,150 mg/l). Absorption maxima (λmax) for RTB H5G was calculated by dissolving a known amount of dye (1mg/ml) and recording its ΔOD from 200nm-800nm. The λmax for RTB H5G was confirmed at 518nm. Amongst all the dye concentrations tried, concentrations 10mg/l, 30mg/l, 50mg/l and 70mg/l showed decolourisation between 85% - 94% within 6-12 days (Table 3, Fig. 1). While the concentrations 50mg/l and 70mg/l showed more than 50% decolourisation. Lowest decolourisation percentage (15.62%) was obtained in 150mg/l dye concentration in 21 days (Table 3, Fig. 1). Variation in the time required and efficiency of decolourisation may probably be due to molecular complexity of dyes and the enzymes produced during decolourisation (Sanghi *et al*., 2006). Slower rate of decolourisation higher is molecular weight and presence of inhibitory groups like nitro (–NO_2_) and sulphite (–SO_3_) in dyes (Hu and Wu, 2001). Maximum decolourisation was observed in shaking conditions as compared to static conditions. This could be due to better oxygen transfer and nutrient distribution as compared to stationary cultures (Kaushik *et al*., 2009). Hence, subsequent dye decolourisation experiments were carried out under shaking condition.

**Fig. 1.**
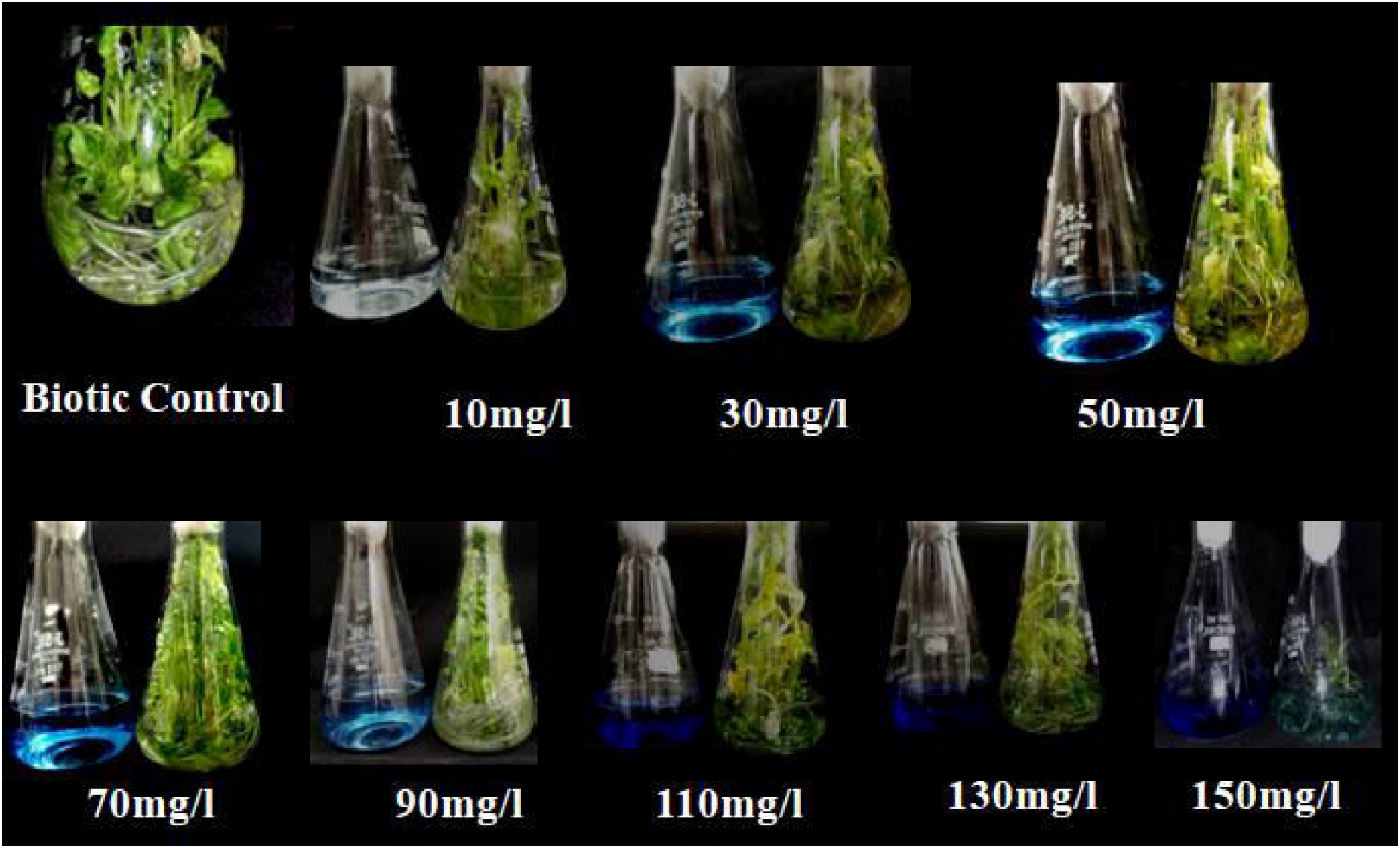
Decolourisation of Reactive Turquoise Blue H5G by *in vitro* plantlets of *Solanum virginianum*

### 4.2 Biochemical analysis of *in vitro* untreated plant of *S. virginianum*

Biochemical analysis and antioxidant potential of plant is determined to verify its medicinal properties. This investigation deals with a remedy for reclamation of polluted sites due to textile dyes. Preliminary studies on decolourisation potential of *S. virginianum* was tested. Medicinal constituents of the plant (TPC, TFC, TAC) and antioxidant capacity (FRAP,FICA,SOAS, PMo and DPPH) of treated and untreated plants were evaluated to confirm that plant remains unaltered after treatment with textile dye Reactive Turquoise Blue H5G. This supports the fact that the plant can be safely used for remediation. This type of study is the first report where the plant is evaluated for its medicinal properties before and after treatment.

#### 4.2.1 Total Phenolic Content (TPC)

The total phenolic content of *S. virginianum* was evaluated using extracts of fresh plant material in different solvent systems. The total phenolic content was varying between 10.86±0.26 to 179.58±0.35 mg of Gallic Acid Equivalent/100g of Fresh Weight (Table 4). From the results it was observed that, lowest activity was found in acetone extract and the highest phenolic content was found in distilled water extract. From the above results it is revealed that, ethanolic and aqueous extracts were beneficial for the extraction of chemical constituents compared to other solvents (Gavande *et al*., 2015). Similar results were obtained in *S. virginianum* by Demla and Varma, 2012 where ethanolic extract was proved to be a better solvent. Phenolic compounds have a wide range of physiological properties such as anti-allergenic, anti-artherogenic, anti-inflammatory, anti-microbial, antioxidant, anti-thrombotic, cardioprotective and vasocardilatory effect (Puupponen *et al*., 2001 and Manach *et al*., 2005).

#### 4.3.2 Total Flavonoid Content (TFC)

The total flavonoid content of *S. virginianum* was evaluated using extracts of fresh plant material in different solvent systems. The total flavonoid content was varying between 0.96±0.01 to 15.76±0.007 mg of Rutin Equivalent/100g of Fresh Weight (Table 4). From the results it was observed that, lowest flavonoid content was found in n-hexane extract and the highest flavonoid content was found in ethanolic extract. Similar results were found in *S. virginianum* where ethanolic extracts shows better results of flavonoid content (Demla and Varma, 2012). Flavonoids are potent antioxidants and capable of scavenging free radicals. Many have antiallergic, antiviral actions and some of them provide protection against cardiovascular mortality (Hertog *et al*., 1993).

#### 4.3.3 Total Alkaloid content

The total alkaloid content of *S. virginianum* was evaluated using extracts of fresh plant material in different solvent systems. The total alkaloid content varies between 39.16±0.05 to 64.12±0.11 mg of Quercitin Equivalent/100g of Fresh Weight (Table 4). From the results it was found that, the lowest activity was found in n-hexane extract and the highest alkaloid content was obtained from ethanolic extract. Alkaloids are a special group of chemicals that are active at different cellular levels of organisms. Alkaloids are detected in about 20% plant species (Sreevidya, 2003). The potent biological activity of some alkaloid had led to their exploitation as pharmaceuticals, narcotics and poisons (Facchini, 2001). Steroidal alkaloid solasodine is the principal alkaloid. Alcoholic extracts of the plant contain fatty and resinous substances. Solasonine is present in fruits. Fruits contain solasonine, solamargine, solanocarpine, beta-solamargine and solanocarpidine. Dry fruits are consisting of traces of isochlorogenic, neochronogenic, chronogenic and caffeic acids. Petals has apigenin whereas stamens has quercetin diglycoside and sitosterol (Kushwaha and Narayan; 2018). Tupkari *et*.*al*., 1972 reported the presence of coumarins, scopolin, scopoletin, esculin and esculetin from plant parts of *S. xantocarpum*.

#### 4.3.4 Antioxidant analysis

The antioxidant potential of the plant extract of *S. virginianum* was measured by different antioxidant assay like, ferric reducing antioxidant power assay(FRAP), ferrous ion chelating assay(FICA), superoxide anion scavenging assay(SOAS), phosphomolybdenum reducing power assay(PMo), 2,2-diphenyl-1-picrylhydrazyl (DPPH) radical scavenging assay. The results of antioxidant potential of the fresh plant material varied according to the nature of solvent used.

In the FRAP activity (Table 4), it was observed that, methanolic extract exhibit highest activity (78.78±0.03mg Ascorbic Acid Equivalent/100g Fresh Weight) while the lowest activity was observed in chloroform extract (70.87±0.47mg AAE/100g FW). In the FICA activity (Table 4), it was found that, acetone extract shows highest activity (30.71%) whereas; lowest activity was seen in distilled water extract (1.35%). In the SOAS activity (Table 4), it was observed that, ethanol extract showed highest activity (46.22%) and lowest activity was observed in distilled water extract (3.25%). In the PMo activity (Table 4), highest activity was observed in acetone extract (66.99%) while methanol extract shows lowest activity (22.79%). In the DPPH activity (Table 4), it was found that, methanol extracts exhibits highest activity (34.38%) while distilled water extract exhibits lowest activity (9.68%).

### 4.4 Biochemical and antioxidant analysis of biotic control plantlets of S. virginianum (treated with Reactive Turquoise Blue H5G dye)

Total Phenolic Content, Total Flavonoid Content, Total Alkaloid Content and Antioxidant ability of *S. virginianum* was studied when it was treated with different concentrations of Reactive Turquoise Blue H5G dye like, 10mg/l, 30mg/l, 50mg/l, 70mg/l, 90mg/l, 110mg/l, 130mg/l and 150mg/l.

#### 4.4.1 Total Phenolic Content (TPC)

The total phenolic content of *S. virginianum* was evaluated using extracts of *in vitro* grown plantlets treated with different concentrations of the dye RTB H5G in different solvent systems. In 10mg/l, the total phenolic content was varying between 9.89±0.02 to 21.48±0.07 mg of GAE/100g of FW (Table 5). From the results it was observed that, the lowest content was found in distilled water extracts and the highest phenolic content was found in chloroform extract. In 30mg/l, the total phenolic content was varying from 95.73±0.02 to 132.91±0.02 mg of GAE/100g of FW (Table 5). From the results it was found that, n-hexane extracts showed lowest content and distilled water extracts showed highest content. In 50mg/l, the total phenolic content was varying between 55.48±0.11 to 78.55±0.14 mg of GAE/100g of FW (Table 5). From the results it was found that, lowest content was observed in n-hexane extracts and the highest content was observed in acetone extracts. In 70mg/l, the total phenolic content was varying from 60.09±0.21 to 115.48±0.50 mg of GAE/100g of FW (Table 5). From the results it was found that, the lowest content was found in distilled water extracts and methanol extracts while the highest content was observed in ethanol extracts. In 90mg/l, the total phenolic content was varying from 65.73±0.25 to 200.35±0.58 mg of GAE/100g of FW (Table 5). From the results it was found that, methanol extracts shows lowest phenolics whereas distilled water extracts shows highest phenolics. In 110mg/l, the total phenolic content was varying from 16.76±0.17 to 55.48±0.51 mg of GAE/100g of FW (Table 5). From the results it was observed that, the lowest content was found in acetone extracts and the highest phenolics were found in ethanol extracts. In 130mg/l, the total phenolic content was varying from 20.86±0.17 to 36.76±0.32 mg of GAE/100g of FW (Table 5). From the results it was observed that, methanol extracts shows lowest phenolics while chloroform extracts shows highest phenolics. In 150mg/l, the total phenolic content was varying from 217.53±0.20 to 364.45±0.79 mg of GAE/100g of FW (Table 5). From the results it was observed that, lowest content was observed in acetone extracts and the highest content was found in chloroform extracts.

#### 4.4.2 Total Flavonoid Content (TFC)

The Flavonoid Content of *S. virginianum* was evaluated using extracts of fresh plant material treated with different concentration of the dye in different solvent systems. In 10mg/l, the total flavonoid content was varying between 6.65±0.04 to11.24±0.14 mg of RE/100g of FW (Table 5). From the results it was observed that, the lowest content was found in n-hexane extract and the highest flavonoid content was found in acetone extract. In 30mg/l, the total flavonoid content was varying between 1.67±0.06 to 9.50±0.27 mg of RE/100g of FW (Table 5). From the results it was observed that, n-hexane extract shows lowest flavonoids while acetone extract shows highest content. In 50mg/l, the total flavonoid content was varying between 1.56±0.06 to 4.07±0.04 mg of RE/100g of FW (Table 5). It was observed that, lowest content was found in distilled water extract while the highest content was found in chloroform extract. In 70mg/l, the total flavonoid content was varying between 0.84±0.09 to 11.81±0.04 mg of RE/100g of FW (Table 5). The results revealed that, lowest content was observed in acetone extract while the highest flavonoids were found in ethanol extract. In 90mg/l, the total flavonoid content was varying between 0.74±0.04 to 7.97±0.01 mg of RE/100g of FW (Table 5). From the results it was observed that, lowest content was observed in acetone extract and the highest content was found in ethanol extract. 110mg/l, the total flavonoid content was varying between 0.94±0.03 to 9.56±0.01 mg of RE/100g of FW (Table 5). From the results it was observed that, n-hexane extract shows lowest flavonoids whereas methanolic extracts shows highest flavonoids. 130mg/l, the total flavonoid content was varying between 0.59±0.04 to 3.73±0.04 mg of RE/100g of FW (Table 5). From the results it was observed that, n-hexane extract shows lowest content while distilled water extract shows highest content. In 150mg/l, the total flavonoid content was varying between 0.61±0.32 to 53.75±0.02 mg of RE/100g of FW (Table 5). From the results it was observed that, ethanol extract shows lowest flavonoids and the chloroform extract shows highest flavonoids.

#### 4.4.3 Total Alkaloid Content

The Alkaloid Content of *S. virginianum* was evaluated using extracts of fresh plant material treated with different concentration of the dye in different solvent systems. In 10mg/l, the total alkaloid ranges between 36.10±0.04 to 60.90±0.17 mg of QE/100g of FW (Table 5). From the results it was found that, n hexane extracts show lowest results whereas acetonic extracts shows highest. In 30mg/l it was found that lowest content was observed in n-hexane extracts and distilled water extracts shows highest alkaloid content and it varies between 25.76±0.15 to 54.55±0.33 mg of QE/100g of FW (Table 5). In 50mg/l, the alkaloid content varying between 30.49±0.06 to 58.95±0.51 mg of QE/100g of FW (Table 5), lowest content was observed in n-hexane extracts and the highest results were observed in methanol extracts. In 70mg/l, it was observed that, distilled water extracts show lowest alkaloid content while the methanolic extracts shows highest alkaloid content and it varies between 6.05±0.06 to 57.16±0.06 mg of QE/100g of FW (Table 5). In 90mg/l, it was seen that, lowest alkaloids were seen in n-hexane extracts whereas methanolic extract shows highest alkaloid content, it ranges between 39.18±0.01 to 55.51±0.23 mg of QE/100g of FW (Table 5). In 110mg/l, the alkaloid content ranges from 25.73±0.02 to 51.40±0.18 mg of QE/100g of FW (Table 5), the lowest results were observed in chloroform extracts and the highest results were obtained from distilled water extracts. In 130mg/l, it was observed that, n-hexane extracts show lowest alkaloid content while the distilled water extract shows highest results, it varies from 25.55±0.01 to 52.49±0.08 mg of QE/100g of FW (Table 5). In 150mg/l, the alkaloid content varies from 21.47±0.02 to 52.94±0.24 mg of QE/100g of FW (Table 5), the lowest results were seen in n-hexane extracts whereas the highest results were observed in methanol extracts.

In the present study fluctuations in the phenolic, flavonoid and alkaloid content was seen with different concentration of dye and with the different solvents used for extraction. This may be due to the stress experienced by the plant because of the dye concentrations.

### 4.4.4 Antioxidant analysis

The plant treated with different concentration of Turquoise Blue H5G dye that is in 10mg/l, the FRAP activity was seen highest in ethanolic extract (44.31±0.03 mg AAE/100g FW) (Table 5) while lowest in chloroform extract (22.82±0.06 mg AAE/100g FW) (Table 5). The FICA activity (Table 5) was found highest in distilled water extract (98.19%) and lowest in chloroform extract (23.05%). The SOAS activity (Table 5) was observed highest in distilled water extract (87.45%) and lowest in chloroform extract (22.30%). The PMo activity (Table 5) was found highest in distilled water extract (95.99%) and lowest in chloroform extract (66.75%), the DPPH activity (Table 5) was found highest in Ethanol extract (14.29%) while lowest was observed in acetone extract (11.38%). In 30mg/l, the FRAP activity was observed highest in distilled water extract (23.01±0.02mg AAE/100g FW) (Table 5) and lowest in n-hexane extract (19.14±0.08mg AAE/100g FW) (Table 5). The FICA activity (Table 5), acetone extract shows highest (86.08%) while ethanol extract shows lowest activity (15.36%). The SOAS activity (Table 5) was observed highest in distilled water extract (60.48%) and lowest in chloroform extract (26.45%). The PMo activity (Table 5) was observed highest in methanol extract (86.88%) and lowest in chloroform extract (57.44%), DPPH activity (Table 5) was highest in ethanol extract (90.10%) and lowest in n-hexane extract (88.92%). In 50mg/l the FRAP activity was found highest in ethanolic extract (46.73±0.04mg AAE/100g FW) (Table 5) and lowest in acetone extract (39.66±0.08mg AAE/100g FW) (Table 4). The FICA activity (Table 5) was found highest in distilled water extract (98.15%) and lowest in chloroform extract (21.96%). In the SOAS activity (Table 5), it was observed that methanolic extract shows highest (79.88%) and lowest in acetone extract (37.22%). In the PMo activity (Table 5), highest activity was observed in n-hexane extract (93.27%) and lowest in distilled water extract (44.27%), the DPPH activity (Table 5) was seen highest in acetone extract (88.71%) and lowest in ethanol and chloroform extract (70.26%). In 70mg/l the FRAP activity was observed highest in methanolic extract (8.60±0.05mg AAE/100g FW) (Table 4) while lowest in distilled water extract (3.70±0.04mg AAE/100g FW) (Table 5). The FICA activity (Table 5) was observed highest in ethanol extract (63.68%) and lowest in chloroform extract (19%). In the SOAS activity (Table 5), the highest activity was found in methanol extract (78.70%) and lowest was observed in n-hexane extract (3.36%). In the PMo activity (Table 5), highest activity was found in acetone extract (93.35%) and lowest was found in distilled water extract (51.96%), DPPH activity (Table 5) was observed highest in methanol and ethanol extract (33.02%) and lowest in acetone extract (29.68%). In 90mg/l the FRAP activity was observed highest in methanol extract (13.74±0.08mg AAE/100g FW) (Table 5) and the lowest activity was found in distilled water extract (1.79±0.07mg AAE/100g FW) (Table 5). The FICA activity (Table 5) was seen highest in methanol extract (92.97%) and lowest in distilled water extract (29.85%). In the SOAS activity (Table 5), highest was observed in chloroform extract (78.22%) and the lowest activity was observed in distilled water extract (12.44%). In the PMo activity (Table 5), highest was found in chloroform extract (91.11%) and lowest activity was observed in distilled water extract (54.73%), the DPPH activity (Table 5) was found highest in methanol extract (30.16%) while chloroform showed lowest activity extract (14.15%). In 110mg/l the FRAP activity was seen highest in n-hexane extract (67.81±0.67mg AAE/100g FW) (Table 5) and lowest activity was seen in ethanol extract (48.64±0.07mg AAE/100g FW) (Table 5). The FICA activity (Table 5) was highest in methanol extract (67.25%) and lowest in chloroform extract (11.54%). In the SOAS activity (Table 5), it was observed highest in acetone extract (87.68%) and lowest in distilled water extract (6.28%). In the PMo activity (Table 5), highest activity was found in n-hexane extract (84.49%) and lowest in chloroform extract (16.54%), DPPH activity (Table 5) was highest seen in distilled water extract (14.65%) and lowest in ethanol extract (4.99%). In 130mg/l the FRAP activity was observed highest in ethanol extract (69.48±0.03mg AAE/100g FW) (Table 5) and lowest in n-hexane extract (26.14±0.10mg AAE/100g FW) (Table 5). The FICA activity (Table 5) was highest in ethanol extract (65.08%) and lowest in chloroform extract (24.66%). The SOAS activity (Table 5) was found highest in n-hexane extract (59.12%) and lowest in ethanol extract (26.58%). The PMo activity (Table 5) was observed highest in chloroform extract (86.45%) and lowest in distilled water extract (57.21%), the DPPH activity (Table 5) was found highest in chloroform extract (18.44%) and lowest in n-hexane extract (3.10%). In 150mg/l the FRAP activity was found highest in ethanol extract (5.73±0.01mg AAE/100g FW) (Table 5) and lowest in methanol extract (4.29±0.02mg AAE/100g FW) (Table 5). The FICA activity (Table 5) was found highest in methanol extract (78.90%) and lowest in chloroform extract (34.15%). The SOAS activity (Table 5) was observed highest in acetone extract (54.03%) and lowest in distilled water extract (17.70%). The PMo activity (Table 5) was found highest in ethanol extract (62.55%) and lowest in acetone extract (28.27%), DPPH activity (Table 5) was observed highest in methanol extract (43.88%) and lowest in n-hexane extract (10.88%). The present study indicates that the plant does not lose their potential even after treated with textile dyes and hence the plant can be successfully used in phytoremediation.

## 5. Phytotoxicity study

Phytotoxicity study was performed to assess the toxicity of dye effluent on common agricultural crop *Vigna radiata*. Untreated dye effluent have caused serious environmental and health problems. Despite this fact they are being released in the water bodies which are used for agricultural purpose. This has direct impact on soil fertility and agricultural productivity. It is relevant to assess the phytotoxicity of the dyes before and after degradation, as environmental safety demands both pollutant removal and their detoxification (Coste, Reynier, 2010; Sartale *et al*., 2010; Dhanve *et al*., 2008). In the present study phytotoxicity was tested on *Vigna radiata* (Mung bean). Percentage of seed germination, shoot length, root length and fresh weight was studied in relation to degraded dye.

In the present study, the untreated synthetic solution of dye in MS medium inhibited the seed germination, shoot length, root length and fresh weight of plants of *Vigna radiata* compared to the degraded metabolite product obtained by individual plants of *Vigna radiata* after decolourisation. The RTB H5G dye solution (untreated sample) showed inhibition of germination, reduced the shoot length and root length as well as affected the fresh weight of plant simultaneously as compared to the degraded metabolite product obtained after decolourisation (Table 6, Fig. 2a).

**Fig. 2a.**
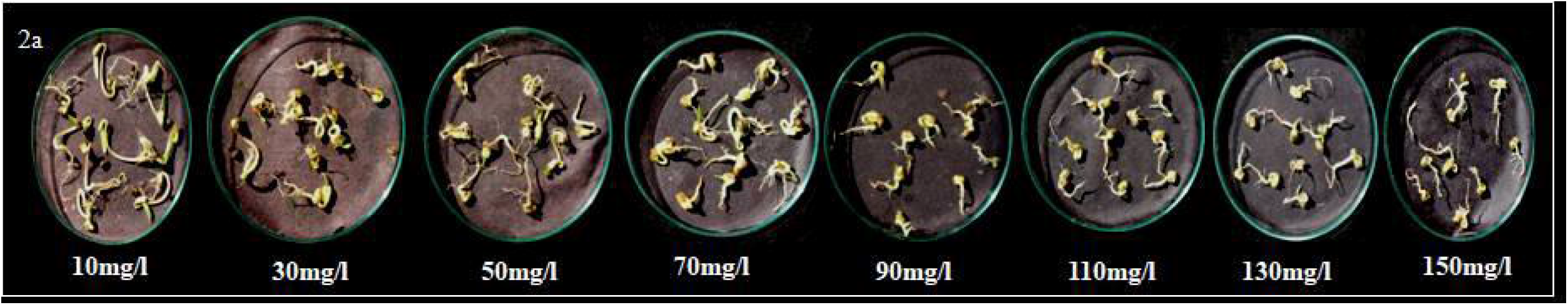
Phytotoxicity studies in *Vigna radiata* seeds germinated in Reactive Turquoise Blue Dye

The degraded products from all the treatment flasks was collected separately and used for watering the seeds of *Vigna radiata*. Ten seeds per treatment were placed in petriplates with blotting papers and the lethality of degraded solution was tested by watering the seeds regularly with the respective degraded products. The germination results were noted down (Table 7, Fig. 2c) and it was observed that the degraded solution 10mg/l, 30mg/l, 50mg/l, 70mg/l, 90mg/l and 110mg/l showed 100% seed germination. Degradation product collected from 130mg/l and 150mg/l treatments showed lowered germination as compared to control (Fig. 2b) treatment. The plumule length, radical length and fresh weight were found to decrease as compared to control treatment. This indicates that though the plant growth was affected by higher treatments, the plant still had potential of decolourising. The phenolics, flavonoids and alkaloids together contribute towards the medicinal properties of plants. The studies of TPC, TFC, TAC (in plants before and after treatment) reveal that the content is not decreased in the treated plantlets. This confirmed that medicinal property of plant is retained even after dye treatment. The overall phytotoxicity study indicated that there was a negative influence of dye and a positive influence regarding non-toxicity of degraded products after decolourisation.

**Fig. 2b.**
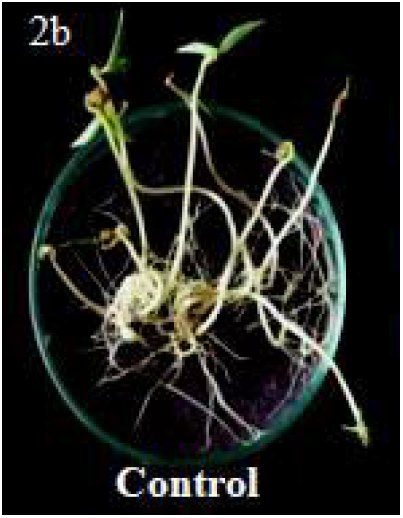
Phytotoxicity studies in *Vigna radiata* seeds germinated in Distilled water. (Control)

**Fig. 2c.**
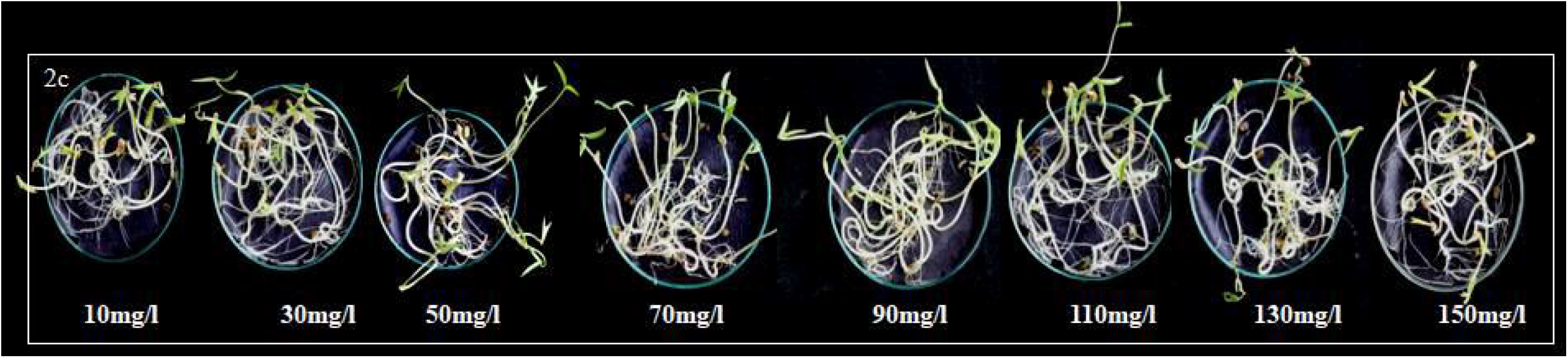
Phytotoxicity studies in *Vigna radiata* seeds germinated in degraded metabolites after decolourisation

## 6. DISCUSSION

*In vitro* studies have been reported in *S. virginianum, S. nigrum, Withania somnifera, Datura metal* and other Solanaceae members. (Sundari *et al*., 2010; Yadav *et al*., 2010; Ramar and Nandagopalan, 2011; Sundar *et al*.,2011; Kumari *et al*., 2013 and Dalavi and Patil, 2017). Selection of explants and hormone addition plays crucial role in plant regeneration. Highest shoot multiplication was reported in *S. virginianum* (Rahman *et al*., 2011). BAP has been more effective for shoot regeneration than Kn in *S. virginianum* (Sundar *et al*., 2011). Present study is in correspondence with Sundar *et al*., (2011) and Rahman *et al*., (2011), as BAP showed best response for shoot regeneration and shoot elongation, when MS medium supplemented with 0.75mg/l BAP. Promising effect of BAP was reported in *Santalum album* (Mujib, 2005) campared to Kn on direct adventitious shoot regeneration. BAP or Kn showed similar results on *in vitro* shoot production of *Solanum nigrum* (Bhat *et al*., 2010) Pawar *et al*., 2002 achieved similar kind of results in *Solanum surattense*. The *in vitro* grown plantlets approximately 4-5cm was used for decolourisation of textile dye Reactive Turquoise Blue H5G. *In vitro* plants were used to maintain the aseptic conditions for growth and to avoid the contamination and plant microbe interactions (Kagalkar *et al*., 2009).

Currently many studies indicate that not only exoenzymes extracted by plant roots and the microoraganisms enzymes are important for the degradation of xenobiotics but also that this may take place in the plant body through its own intrinsic enzymatic machinery (Araujo *et al*. 2002). The existence of plant tolerant to high concentrations of phenolic compounds suggests their ability to degrade those compounds via peroxidase (POD) (Naghibi *et al*. 2003). Plants are unique having remarkable metabolic and absorptional skills, along with transport system. Following uptake, the main purpose is to convert the lipophilic xenobiotics into a more water soluble and less toxic metabolite that can therefore be eliminated (Alkorta and Garbisu 2001). Plant detoxification pathway implicates specific enzymes in three distinct phases. Oxidases or oxigenases or carotenoid monoxygenases and dioxygenases might be involved in the first transformation step. Phase 2 involves the xenobiotic conjugation by glutathione -s- transferase (GST). Finally, in phase 3, plant internal screening-off area and elimination of the metabolites formed can be accomplished by storage in the cell vacuole or co-valent binding to cell walls (Coleman *et al*. 1997; Burken *et al*.,2004 Schroder *et al*. 2007; Schwitzuebel *et al*., 2011). Phytoremediation takes the advantage of the unique and selective uptake capabilities of plant root system, together with the translocation, bioaccumulation and contaminant degradation abilities of the entire plant body. Accumulators survive despite concentrating contaminants in their aerial tissues, they biodegrade or biotransform the contaminants into inert form in their tissues. The excluders restrict contaminant uptake into their biomass. Plants have exceptional potential to concentrate and accumulate elements and compounds in addition to this it can also uptake contaminants from the environment and they have evolved an advanced regulatory mechanism to metabolize them into various molecules in their organs and tissues (Salt *et al*., 1998). Plants help to clean up various pollutions that include metals, pesticides, explosives and oil, also help to prevent carrying of pollutants and products of contaminants away from sites to other areas by wind, rain and groundwater (Cluis, 2004). Phytoremediation system requires less amount of nutrient input (Cunnighum and Berti, 2000).

Recent studies using plants for reclamation have confirmed their textile dye-degradation ability. These mainly included plants like *Tagetes patula, Glandularia pulchella, Aster amellus, Petunia grandiflora, Salsola vermiculat, Sesuvium portulacastrum* etc. (Bestani *et al*., 2008; Khandare *et al*., 2011a, 2011b; Kabra *et al*., 2011a, 2011b; Patil *et al*., 2012; Watharkar *et al*., 2013b). Remediation of textile dyes like Acid Orange 7, Direct Red 5B, Brilliant Blue R, Remazol Red, sulphonated anthraquinone and acid dye has been independently conducted by using some plants namely *Thymus valgaris, Tagetes patula, Rheum rabarbarum, Rheum hydroplantarum and Brassica rapa* (Zheng and Shetty, 2000; Aubert and Schwitzguebel, 2004; Kulshrestha and Husain, 2007; Patil *et al*., 2009). Similar experiment was carried out with *in vitro* grown plantlets of *Petunia grandiflora, Gailardia grandiflora* and their consortium for decolourisation of dyes Brilliant Blue G, Direct Blue GLL, Rubin GFL, Scarlet RR and Brown 3 REL. Brown 3 REL was decolourised up to 81, 79 and 91% by *P. grandiflora, G. grandiflora* and their consortium respectively within 36h. Direct Blue GLL decolorized up to 78, 72 and 91% within 48h (Watharkar and Jadhav, 2014). *P. grandiflora* showed versatility in decolourising Brilliant Blue G was decolourised up to 86 % without any adsorption within 36h by wild plants as well as tissue cultures of *P. grandiflora* (Watharkar *et al*., 2013).

Plants have an innate ability to synthesize an array of antioxidants capable of arresting the ROS (Reactive Oxygen Species) causing oxidative damage (Kasote *et al*., 2015). Plant antioxidants are chemicals like phenolics, flavonoids which have specific roles in phytochemical responses towards stress. Using more than a single method is therefore suggested to understand the exhaustive and complete prediction of antioxidant potential from the actual collected data (Luximon-Ramma *et al*., 2002). The present study was focussed on antioxidant capacity of biotic control and the plant treated with the textile dye. Mechanism of antioxidant action can include suppression of ROS formation by inhibition of enzymes or by chelating trace elements involved in free radical generation. Phenolic compounds are classified into phenolic acids, flavonoid polyphenolics (flavones, flavonones, xanthones and catechins) and non-flavonoid polyphenolics. Phenolic compounds act as free radical terminators (Shahidi *et al*., 1992). Due to presence of hydroxyl group which is efficient of scavenging the free radicals, phenolics and antioxidant activity are positively correlated (Vinson *et al*., 1998). Phenolic compound donates electron to scavenge hydrogen peroxide and convert it into water (Nabavi *et al*., 2009). Presence of sugar and hydroxyl group makes flavonoid a water soluble whereas isopentyl units and methyl group makes them lipophilic (Crozier *et al*., 2006). Flavonoid donates hydrogen atom to free radical so immediately that it interferes in further oxidation of lipids and other molecules (Schroeter *et al*., 2002). The studies have reported the role of flavonoids in secondary antioxidant defence mechanism in stress exposed plant (Agati *et al*., 2012). The total phenolic content and total flavonoid content was studies successfully in *S. virginianum* by Morankar *et al*., (2019) in methanolic extract. Kushwaha and Narayan (2018), studied total phenolics and total flavonoids in *S. xanthocarpum* in methanolic extracts. The most common method of extending the application of antioxidants activity of plants is DPPH which is commercial, stable, common and organic free radical (Gadade and Patil; 2019). DPPH free radical scavenging is an accepted mechanism for screening the antioxidant activity of plant extracts. DPPH assay detects antioxidant such as flavonoids, polyphenols. Ferrous ion catalysing oxidation exerts tremendous stress on plant cells. To make it less severe an effective and common food pro-oxidant, ferrous ion was used. The transition metal ferrous Fe^2+^ is trapped in the antioxidant of the plant extracts. The brick red colour of ferrozine Fe^2+^ complex is reduced. Phenazine methosulfate-nicotinamide adenine dinucleotide (PMS/NADH) system which generates superoxide radicals, reduce nitro blue tetrazolium (NBT) to a purple formazan. The active compounds in the plant extract inhibit the reduction of NBT by quenching the superoxide radicals. Phosphomolybdenum reducing power assay is based on the reduction of Mo (VI) to Mo (V) by sample analyte and subsequent formation of a green phosphate-Mo (V) complex at acidic pH, usually detects antioxidants such as some phenolics, ascorbic acid, α – tocopherol and carotenoids (Diwan *et al*., 2012). The FRAP assay measures the antioxidant potential in samples through the reduction of ferric tripyridyltriazine (Fe^3+^ - TPTZ) complex to ferrous (Fe^2+^ - TPTZ) at low pH by the reductant compounds present in the plant extract indicating antioxidant potential. The phytochemical screening of the plants revealed the presence of compounds like Phenols, flavonoids, saponins, terpenoids, steroids, and glycosides. They act as allelochemicals, released in the surrounding environment (Gagare *et al*., 2021). The present studies thus revealed that the medicinal properties of *S. virginianum* were evaluated in terms of total phenolics, flavonoids, alkaloids which in turn support the antioxidant nature of *S. virginianum*, this investigation is a pre-requisite protocol for determination of loss of medicinal properties of *S. virginianum* when treated with RTB H5G dye. The studies were carried out preliminarly to check the medicinal status of *S. Virginianum* plants treated with RTB H5G dye (one-time addition of RTB H5G dye). The studies may further be designed for repetitive use of same plantlet while there is continuous and periodic addition of the respective dye. The experiment may help to monitor the exact time of replacement of plants used for phytoremediation.

Phytotoxicity study on *Phaseolus mungo* and *Sorghum vulgare* was done successfully in treated dye solution using *Gailardia grandiflora, Petunia grandiflora* and their consortium (Watharkar and Jadhav, 2014). The lower percentage of germination of *Phaseolus mungo* and *Sorghum vulgare* with lower length of plumule and radical in the dye solution indicate that the dye, Green HE4B was toxic to these plants, while the metabolites formed after the degradation of the dye by *Glandularia pulchella* were almost as non-toxic as distilled water (Kabra *et al*., 2011). Similar study was carried out successfully in a degradation of a disperse, disulfonated triphenylmethane textile dye Brilliant Blue G in *Petunia grandiflora* (Watharkar *et al*., 2013).

## 7. CONCLUSION

*S. virginianum* (L.) decolourise textile dye Reactive Turquoise Blue H5G which reveals that it is the best material for reclamation of textile dye polluted sites. The fate of degraded metabolites is however not known. Degraded metabolites have a non-poisonous nature which was confirmed by phytotoxicity studies. Biochemicals and Antioxidant potential of the plant retains throughout the treatment which signifies the potent use of plants for remediation of pollution sites proving the technology to be safer.

## ACKNOWLEDGEMENT

Author 1 is grateful to the Department of Botany, Shivaji University Kolhapur for providing laboratory facilities.

