## Supplementary material for "PHYTOREMEDIATION OF REACTIVE TURQUOISE BLUE H5G USING *IN VITRO* CULTURES OF *SOLANUM VIRGINIANUM* (L.)": Tables

Table 1 *In vitro* seed germination of *S. virginianum*

| **Sr. No.** | **Medium** | **No. of seeds inoculated** | **No. of seeds germinated** | **% of seed germination** |
| --- | --- | --- | --- | --- |
| 1. | ½ MS Basal | 300 | 197 | 65.66 |

MS basal medium containing ½ strength major nutrients, minor nutrients, vitamins, Fe EDTA, inositol, sucrose 1.5% and agar 0.8%.

Table 2 *In vitro* shoot multiplication in *S.* *virginianum*

| **Sr. No.** | **Concentration of BAP (mg/l)** | **Mean no. of shoots ± SE** | **% response** |
| --- | --- | --- | --- |
| 1. | 0.5 | 12.8±0.28 | 80 |
| 2. | 0.75 | 18.1±0.17 | 100 |
| 3. | 1.0 | 14.1±0.22 | 40 |

Values represent mean ± S. E. of twenty replicates per treatment and all the experiments were repeated thrice.

Table 3	**Decolourisation** **of textile dye RTB H5G** by *in vitro* plantlets of *S*. *virginianum*

| **Concentration of the dye**  **mg/l** | **Decolourisation**  **(%)** | **Days required for maximum decolourisation** |
| --- | --- | --- |
| **10** | 93.98 | 6 |
| **30** | 89.36 | 8 |
| **50** | 89.20 | 8 |
| **70** | 85.36 | 12 |
| **90** | 60.00 | 15 |
| **110** | 52.12 | 19 |
| **130** | 44.44 | 21 |
| **150** | 15.62 | 21 |
| **Biotic control(devoid of dye)** | --- | --- |

Approximately 1g of *in vitro* grown *S. virginianum* plantlets were suspended in each treatment flask.

**Table 4:** **Biochemical and antioxidant analysis of biotic control plantlets of S. virginianum (untreated with Reactive Turquoise Blue H5G dye)**

| Sr. No. | Solvent | TPC | TFC | TAC | FRAP | FICA  (%) | SOAS  (%) | PMo  (%) | DPPH  (%) |
| --- | --- | --- | --- | --- | --- | --- | --- | --- | --- |
| 1. | Methanol | 87.53±0.11 | 10.41±0.03 | 44.39±0.08 | 78.78±0.03 | 7.09±0.01 | 30.85±0.01 | 22.79±0.07 | 34.38±0.002 |
| 2. | DW | 179.58±0.35 | 10.01±0.01 | 50.83±0.06 | 77.05±0.39 | 1.35±0.01 | 3.25±0.02 | 38.08±0.01 | 9 9.68±0.002 |
| 3. | Ethanol | 135.99±0.26 | 15.76±0.07 | 64.12±0.11 | 73.23±0.09 | 16.66±0.02 | 46.22±0.04 | 28.71±0.05 | 32.65±0.005 |
| 4. | Chloroform | 26.24±0.20 | 4.24±0.03 | 41.11±0.11 | 70.87±0.47 | 25.11±0.04 | 30.98±0.09 | 52.30±0.03 | 21.27±0.005 |
| 5. | Acetone | 10.86±0.26 | 2.70±0.01 | 56.03±0.12 | 72.26±0.16 | 30.71±0.05 | 43.88±0.08 | 66.99±0.07 | 11.07±0.001 |
| 6. | n-hexane | 27.01±0.29 | 0.96±0.01 | 39.16±0.05 | 77.81±0.62 | 25.33±0.03 | 44.40±0.08 | 62.37±0.01 | 10.90±0.003 |

Values represent mean of triplicate reading ± SE.

*mg GAE/100g FW – milligrams Gallic Acid Equivalents per 100 grams Fresh Weight.

**mg RE/100g FW - milligrams Rutin Equivalents per 100 grams Fresh Weight.

***mg QE/100g FW - milligrams Quercitin Equivalents per 100 grams Fresh Weight.

****mg AAE/100g FW - milligrams Ascorbic Acid Equivalents per 100 grams Fresh Weight.

**Table 5:** **Biochemical and antioxidant analysis of biotic control plantlets of S. virginianum (treated with Reactive Turquoise Blue H5G dye)**

| Sr.No | Conc. of dye | Solvent | TPC* | TFC** | TAC*** | FRAP**** | FICA  (%) | SOAS  (%) | PMo  (%) | DPPH  (%) |
| --- | --- | --- | --- | --- | --- | --- | --- | --- | --- | --- |
| 1. | 10mg/l | Methanol | 20.29±0.09 | 7.05±0.04 | 45.89±0.32 | 44.25±0.08 | 96.76±0.004 | 58.36±0.002 | 94.94±0.01 | 13.46±0.002 |
|  |  | DW | 9.89±0.02 | 7.26±0.03 | 50.79±0.37 | 38.66±0.27 | 98.19±0.006 | 87.45±0.005 | 95.99±0.002 | 12.18±0.009 |
|  |  | Ethanol | 16.48±0.15 | 8.42±0.05 | 52.76±0.31 | 44.31±0.03 | 96.60±0.003 | 54.84±0.01 | 93.94±0.005 | 14.29±0.006 |
|  |  | Chloroform | 21.48±0.07 | 7.66±0.13 | 48.23±0.10 | 22.82±0.06 | 23.05±0.001 | 22.30±0.01 | 66.75±0.008 | 12.32±0.001 |
|  |  | Acetone | 19.02±0.22 | 11.24±0.14 | 60.90±0.17 | 36.57±0.01 | 63.41±0.003 | 22.66±0.003 | 73.59±0.009 | 11.38±0.008 |
|  |  | n Hexane | 17.73±0.20 | 6.65±0.04 | 36.10±0.04 | 33.84±0.01 | 64.13±0.015 | 39.69±0.005 | 92.38±0.01 | 13.08±0.006 |
| 2. | 30mg/l | Methanol | 126.24±0.11 | 2.25±0.16 | 48.79±0.07 | 21.71±0.01 | 68.48±0.009 | 51.98±0.004 | 86.88±0.007 | 89.37±0.006 |
|  |  | DW | 132.91±0.20 | 8.03±0.13 | 54.55±0.33 | 23.01±0.02 | 71.84±0.004 | 60.48±0.003 | 86.71±0.008 | 89.23±0.004 |
|  |  | Ethanol | 107.53±0.09 | 3.99±0.13 | 46.69±0.48 | 21.55±0.01 | 15.36±0.02 | 48.54±0.01 | 82.51±0.002 | 90.10±0.004 |
|  |  | Chloroform | 113.68±0.03 | 3.42±0.31 | 33.17±0.02 | 22.82±0.01 | 60.80±0.001 | 26.45±0.005 | 57.44±0.01 | 89.20±0.009 |
|  |  | Acetone | 102.65±0.10 | 9.50±0.27 | 41.06±0.10 | 21.11±0.03 | 86.08±0.006 | 36.29±0.003 | 75.51±0.01 | 89.71±0.009 |
|  |  | n Hexane | 95.73±0.02 | 1.67±0.06 | 25.76±0.15 | 19.14±0.08 | 56.96±0.002 | 47.68±0.01 | 81.64±0.01 | 88.92±0.003 |
| 3. | 50mg/l | Methanol | 56.5059 | 3.81±0.02 | 58.95±0.51 | 43.53±0.24 | 96.69±0.004 | 79.88±0.01 | 87.24±0.01 | 71.89±0.004 |
|  |  | DW | 59.58282 | 1.56±0.06 | 53.49±0.14 | 42.46±0.05 | 98.15±0.006 | 58.53±0.08 | 44.27±0.009 | 82.27±0.009 |
|  |  | Ethanol | 60.60846 | 2.99±0.03 | 47.77±0.09 | 46.73±0.04 | 96.53±0.003 | 60.83±0.03 | 89.84±0.008 | 70.26±0.004 |
|  |  | Chloroform | 70.35205 | 4.07±0.04 | 31.58±0.009 | 42.26±0.07 | 21.46±0.004 | 48.61±0.005 | 91.53±0.006 | 70.26±0.008 |
|  |  | Acetone | 78.55718 | 3.54±0.01 | 43.33±0.09 | 39.66±0.08 | 62.66±0.005 | 37.22±0.01 | 55.73±0.006 | 88.71±0.003 |
|  |  | n Hexane | 55.48026 | 3.25±0.04 | 30.49±0.06 | 42.79±0.06 | 63.38±0.01 | 37.46±0.01 | 93.27±0.008 | 73.17±0.003 |
| 4. | 70mg/l | Methanol | 60.09±0.25 | 5.62±0.02 | 57.16±0.06 | 8.60±0.05 | 43.99±0.01 | 78.70±0.003 | 74.39±0.009 | 33.02±0.01 |
|  |  | DW | 60.09±0.21 | 2.66±0.02 | 6.05±0.06 | 3.70±0.04 | 33.78±0.01 | 23.93±0.001 | 51.96±0.006 | 29.99±0.001 |
|  |  | Ethanol | 115.48±0.50 | 11.81±0.04 | 48.86±0.11 | 7.30±0.06 | 63.68±0.02 | 69.25±0.03 | 79.24±0.008 | 33.02±0.01 |
|  |  | Chloroform | 64.45±0.05 | 3.94±0.08 | 26.9±0.01 | 4.04±0.38 | 19±0.01 | 56.81±0.008 | 78.80±0.01 | 30.44±0.008 |
|  |  | Acetone | 78.55±0.03 | 0.84±0.09 | 43.60±0.15 | 4.56±0.01 | 19.24±0.01 | 19.77±0.002 | 93.35±0.01 | 29.68±0.006 |
|  |  | n Hexane | 68.81±0.16 | 3.10±0.01 | 27.71±0.11 | 4.00±0.07 | 27.41±0.01 | 3.36±0.02 | 76.04±0.01 | 30.99±0.006 |
| 5. | 90mg/l | Methanol | 65.73±0.25 | 4.00±0.04 | 55.51±0.23 | 13.74±0.08 | 92.97±0.002 | 59.25±0.01 | 87.98±0.01 | 30.16±0.01 |
|  |  | DW | 200.35±0.58 | 1.21±0.01 | 49±0.15 | 1.79±0.07 | 29.85±0.01 | 12.44±0.03 | 54.73±0.01 | 24.10±0.002 |
|  |  | Ethanol | 74.96±0.30 | 7.97±0.01 | 47.64±0.16 | 8.96±0.01 | 78.92±0.002 | 38.12±0.005 | 88.40±0.01 | 25.82±0.01 |
|  |  | Chloroform | 67.53±0.50 | 0.80±0.02 | 41.83±0.01 | 6.92±0.13 | 37.27±0.04 | 78.22±0.004 | 91.11±0.003 | 14.15±0.005 |
|  |  | Acetone | 119.32±0.20 | 0.74±0.04 | 41.42±0.19 | 3.31±0.25 | 35.95±0.01 | 34.81±0.004 | 59.68±0.005 | 26.82±0.009 |
|  |  | n Hexane | 67.78±0.42 | 5.60±0.03 | 39.18±0.01 | 7.29±0.13 | 30.35±0.01 | 18.34±0.01 | 65.36±0.008 | 18.18±0.004 |
| 6. | 110mg/l | Methanol | 35.73±0.06 | 9.56±0.01 | 47.43±0.01 | 56.98±0.21 | 67.25±0.006 | 46.85±0.01 | 83.85±0.009 | 6.13±0.002 |
|  |  | DW | 33.17±0.17 | 4.55±0.01 | 51.40±0.18 | 63.09±0.10 | 36.22±0.008 | 6.28±0.01 | 77.10±0.007 | 14.65±0.01 |
|  |  | Ethanol | 55.48±0.51 | 1.31±0.36 | 48.18±0.09 | 48.64±0.07 | 60.93±0.004 | 51.69±0.006 | 77.75±0.006 | 4.99±0.01 |
|  |  | Chloroform | 23.68±0.04 | 1.16±0.04 | 25.73±0.02 | 63.37±0.67 | 11.54±0.002 | 53.86±0.009 | 16.54±0.002 | 6.06±0.008 |
|  |  | Acetone | 16.76±0.17 | 1.29±0.09 | 40.97±0.08 | 54.34±0.29 | 54.39±0.005 | 87.68±0.01 | 59.43±0.004 | 12.40±0.002 |
|  |  | n Hexane | 35.99±0.11 | 0.94±0.03 | 26.44±0.02 | 67.81±0.23 | 57.05±0.002 | 62.80±0.01 | 84.49±0.007 | 7.10±0.003 |
| 7 | 130mg/l | Methanol | 20.86±0.17 | 1.31±0.05 | 51.76±0.21 | 34.55±0.21 | 64.27±0.01 | 38.88±0.002 | 73.61±0.008 | 4.51±0.002 |
|  |  | DW | 28.30±0.24 | 3.73±0.04 | 52.49±0.08 | 63.85±0.09 | 52.48±0.01 | 54.36±0.005 | 57.21±0.009 | 12.09±0.01 |
|  |  | Ethanol | 23.68±0.17 | 0.90±0.04 | 45.69±0.08 | 69.48±0.03 | 65.08±0.01 | 26.58±0.009 | 70.05±0.01 | 4.55±0.002 |
|  |  | Chloroform | 36.76±0.32 | 3.18±0.04 | 27.03±0.01 | 64.76±0.20 | 24.66±0.03 | 42.46±0.01 | 86.45±0.01 | 18.44±0.003 |
|  |  | Acetone | 22.91±0.08 | 0.84±0.01 | 38.48±0.02 | 26.56±0.29 | 38.54±0.01 | 51.19±0.01 | 77.89±0.01 | 14.06±0.003 |
|  |  | n Hexane | 33.42±0.17 | 0.59±0.04 | 25.55±0.01 | 26.14±0.10 | 49.54±0.01 | 59.12±0.05 | 79.85±0.01 | 3.10±0.005 |
| 8. | 150mg/l | Methanol | 309.83±0.87 | 30.65±0.03 | 52.94±0.24 | 4.29±0.02 | 78.90±0.006 | 46.74±0.006 | 61.01±0.005 | 43.88±0.001 |
|  |  | DW | 264.19±0.17 | 1.45±0.15 | 45.91±0.14 | 5.25±0.02 | 68.45±0.001 | 17.70±0.003 | 40.70±0.003 | 25.64±0.002 |
|  |  | Ethanol | 306.50±0.16 | 0.61±0.32 | 46.78±0.07 | 5.73±0.02 | 41.07±0.003 | 43.35±0.009 | 62.55±0.006 | 14.39±0.005 |
|  |  | Chloroform | 364.45±0.79 | 53.75±0.02 | 27.16±0.02 | 5.33±0.06 | 34.15±0.009 | 52.47±0.003 | 56.04±0.006 | 13.88±0.002 |
|  |  | Acetone | 217.53±0.20 | 40.22±0.05 | 34.80±0.11 | 4.65±0.10 | 77.55±0.001 | 54.03±0.002 | 28.27±0.01 | 13.75±0.008 |
|  |  | n Hexane | 321.37±0.05 | 0.94±0.01 | 21.47±0.02 | 4.74±0.00 | 54.18±0.001 | 24.60±0.01 | 57.32±0.05 | 10.88±0.003 |

Values represent mean of triplicate reading ± SE.

*mg GAE/100g FW – milligrams Gallic Acid Equivalents per 100 grams Fresh Weight.

**mg RE/100g FW - milligrams Rutin Equivalents per 100 grams Fresh Weight.

***mg QE/100g FW - milligrams Quercitin Equivalents per 100 grams Fresh Weight.

****mg AAE/100g FW - milligrams Ascorbic Acid Equivalents per 100 grams Fresh Weight.

**Table 6 Phytotoxicity studies in *Vigna radiata* (watered with MS supplemented with RTB H5G	dye)**

| Parameter | control | 10mg/l | 30mg/l | 50mg/l | 70mg/l | 90mg/l | 110mg/l | 130mg/l | 150mg/l |
| --- | --- | --- | --- | --- | --- | --- | --- | --- | --- |
| % germination | 100 | 100 | 100 | 90 | 90 | 80 | 80 | 60 | 50 |
| Plumule length(cm) | 9.29±0.67 | 4.41±0.41 | 2.62±0.45 | 2.68±0.36 | 2.51±0.24 | 1.82±0.17 | 1.73±0.15 | 1.51±0.10 | 1.41±0.11 |
| Radicle length(cm) | 6.03±0.43 | 2.89±0.21 | 2.72±0.32 | 2.63±0.16 | 1.93±0.27 | 1.82±0.22 | 1.87±0.22 | 1.67±0.15 | 1.51±0.15 |
| Fresh weight(g) | 1.97 | 1.14 | 0.98 | 0.96 | 0.94 | 0.87 | 0.86 | 0.84 | 0.78 |

Readings represents mean value of three replicates per treatment ± SE. Each replicate had ten seeds.

**Table 7 Phytotoxicity studies in *Vigna radiata* (watered with MS supplemented with MS supplemented with degraded product of RTB H5G dye)**

| Parameter | CSontrol | 10mg/l | 30mg/l | 50mg/l | 70mg/l | 90mg/l | 110mg/l | 130mg/l | 150mg/l |
| --- | --- | --- | --- | --- | --- | --- | --- | --- | --- |
| % germination | 100 | 100 | 100 | 100 | 100 | 100 | 100 | 100 | 100 |
| Plumule length (cm) | 9.29±0.67 | 8.93±0.99 | 11.69±0.55 | 14.35±0.94 | 11.43±0.94 | 13.58±0.21 | 12.55±0.80 | 12.97±0.40 | 8.56±0.95 |
| Radicle length (cm) | 6.03±0.43 | 5.45±0.32 | 7.05±0.83 | 7.12±0.69 | 6.45±0.49 | 6.99±0.21 | 7.0±0.50 | 5.97±0.50 | 6.62±0.69 |
| Fresh weight (g) | 1.97 | 2.46 | 2.74 | 2.96 | 2.54 | 2.82 | 2.85 | 2.46 | 2.49 |

Readings represents mean value of three replicates per treatment ± SE. Each replicate had ten seeds.
